# Male rat sexual behavior: insights from inter-copulatory intervals

**DOI:** 10.1101/2021.03.16.435659

**Authors:** Patty T. Huijgens, Fay A. Guarraci, Jocelien D.A. Olivier, Eelke M.S. Snoeren

## Abstract

The assessment of sexual behavior in male rats with the aim of unraveling underlying neurobiological mechanisms has in the recent decades been reduced to the annotation of mounts, intromissions and ejaculations. To provide a better understanding of the structure and patterns of copulation, it is necessary to extend and tailor the analysis to the natural organization of male rat copulation. This will lead to better formulation of hypotheses about neurobiological underpinnings of behavior. Mounts and intromissions are naturally organized in mount bouts consisting of one or more copulatory behaviors and are interspersed with time outs. We hypothesized that time outs and the post-ejaculatory interval (inter-copulatory intervals) are related and possibly under the control of a common copulatory inhibition mechanism that is the result of penile sensory stimulation. To test this hypothesis, we analyzed sexual behavior in male rats of three different cohorts from three different laboratories. Results showed that the post-ejaculatory interval and mean time out duration are strongly correlated in all cohorts analyzed. In addition, we showed that individual time out duration is at least partially predicted by the sum of sensory stimulation of copulatory components in the preceding mount bout, with more penile stimulation associated with longer time outs. These findings suggest that both time out and post-ejaculatory interval duration may be determined by the magnitude of sensory stimulation, which inhibits copulation. Whether the same neural pathways are involved in the central orchestration of both time outs and the post-ejaculatory interval should be subject to future studies.

## 1. Introduction

In order to understand the neurobiological mechanisms underlying the orchestration of copulation, it is important to understand the full range and patterning of the behavior in detail. Recently, a critical perspective has warned against a reductionist bias in behavioral neuroscience and called for more detailed behavioral analysis leading to better foundations for hypothesis generation about neurobiological underpinnings of behavior.^1^ Male rats are an often-used animal model for sexual behavior in both basic and translational neuroscience research. Yet, behavioral annotation of copulation is often limited to the frequency of mounts, intromissions, and ejaculations, including parameters calculated from the times these behaviors occurred (e.g., latencies, post-ejaculatory interval, intromission ratio). A more detailed analysis of the organization and patterns of male rat copulation has been introduced in the past,^2^ but has been underrepresented in studies of the more recent decades.

In the pioneering study by Sachs and Barfield (1970),^2^ it was convincingly demonstrated that male rat copulation is temporally organized in mount bouts, which are defined as “a sequence of mounts (one or more), with or without intromission, uninterrupted by any behavior (other than genital autogrooming) that is not oriented towards the female”. Mount bouts are naturally separated by longer periods of no interaction with the female, defined as “time outs”. This mount bout pattern is not driven by intromissions, as males that can only mount still organize copulation in mount bouts of one or multiple mounts interspersed with time outs. Therefore, the mount bout should be considered the basic unit of copulation, and temporal patterning of copulation (copulatory pace) is better reflected in the time outs between mount bouts than in the more traditionally used inter-intromission interval that disregards mounts.^2^ Copulatory pace is an important pillar of male copulation as it determines the latency to ejaculation together with sensitivity (i.e., number of intromissions needed to reach ejaculation) and efficiency (i.e., achieved intromissions per total mounts). Therefore, pursuing a deeper understanding of the temporal organization of male copulation will contribute to the development of better theoretical concepts of the structure of copulatory behavior.

Like the time out, the post-ejaculatory interval (PEI) could also be considered a parameter of copulatory temporal organization. Both the PEI and the time out are inter-copulatory intervals, be it for different durations (e.g., PEI > time out). It is still unclear what neurobiological mechanisms underly the PEI. It has been shown that the PEI is a result of a central, rather than a peripheral (genital), neuronal inhibition,^3^ and some brain regions and neurotransmitters have been implicated to be involved in the regulation of the PEI (e.g., galanergic signaling in the medial subparafascicular thalamus, and falling levels of glutamate and dopamine in the medial preoptic area; reviewed by Seizert (2018)^4^). But still, the neurobiological orchestration of this strong and partially absolute central inhibition remains to be elucidated. Likewise, the neurobiological regulation of inter-copulatory-intervals that are observed before ejaculation (i.e., time outs), remain elusive. In view of both the PEI and the time out being the result of a copulatory inhibition, both of these inter-copulatory intervals might be regulated by the same neuronal inhibitory mechanism. Therefore, the investigation of how inter-copulatory intervals relate to each other in the complex structure and pattern of male copulatory behavior is important.

Some evidence for the possible relationship between inter-copulatory intervals is found in several correlational and factor analysis studies of male rat sexual behavior.^5–7^ The PEI consistently loads onto the same factor as the inter-intromission interval (III) together with the number of ejaculations and ejaculation latency, referred to as the “copulatory rate factor”. In addition, the PEI, III, and time out are all longer in older compared to younger naive male rats,^8^ and both the PEI and III are shortened upon enforced inter-copulatory intervals (making the female unavailable for a short amount of time),^9^ suggesting a relationship between these parameters. Conversely, the PEI increases over each subsequent ejaculation series, whereas the mean III duration follows a U-shape over ejaculation series.^10^ Following our notion that the time out, and not the III, is the natural inter-copulatory interval before ejaculation, as the mount bout is the basic copulatory unit, we hypothesize that PEI and time out duration are closely related within individual rats, and more strongly correlated than PEI and III.

The PEI is clearly induced by a strong sensory stimulus, namely ejaculation. If the PEI and the time out are related, it is to be expected that time outs are also induced by sensory stimulation in the preceding mount bout. Both mounts and intromissions contribute to achievement of ejaculation, but intromissions provide stronger sensory penile stimulation than mounts.^11^ However, it has been found that prevention of intromissions does not change the distribution of time outs,^2,12^ and the same lab found that the mean time out duration does not depend on the last behavior (mount or intromission) within the preceding mount bout.^13^ Still, intromissions are far more likely to end a mount bout than extravaginal intromissions (motorically identical to intromissions but without penile insertion) or mounts.^13^ These results trigger the question of whether the total sensory stimulation of the sum of copulatory components within the mount bout might predict the duration of the following time out. If so, there would be reason to believe that both ejaculation and mount bout induce a similar copulatory inhibition that is determined by the magnitude of sensory stimulation.

We present a detailed description of the mount bout organization of copulation based on behavioral analysis of three different male rat cohorts from three different laboratories. We assessed correlation of PEI and time out within rats, and how these parameters change over ejaculation series as well as across repeated copulation sessions. Moreover, we determined what mount bout characteristics predict the duration of the directly following time out. Our findings lead us to hypothesize that a central inhibitory mechanism might control both the temporal patterning of copulatory behavior within an ejaculation series, as well as the time in between ejaculation series.

## 2. Materials and Methods

The data presented in this paper consists of three male rat cohorts from three different laboratories in three different locations, from here on referred to as the “Tromsø” (Snoeren lab), “Groningen” (Olivier lab), and “Texas” (Guarraci lab) cohorts.

### 2.1 Animals

#### Tromsø

The data from this cohort comes from a previously published experiment ^14^. For the purpose of this previous experiment, the 53 male Wistar rats (Charles River, Sulzfeld, Germany) of approximately three months old had undergone brain surgery during which a viral construct coding for Designer Receptors Exclusively Activated by Designer Drugs (DREADDs) was infused bilaterally into the medial amygdala. The data set used in the current paper consists of annotations from a copulation test preceded by an intraperitoneal injection with vehicle (deionized water), 45 minutes before the copulation test. Since DREADDs are inert without the ligand clozapine-N oxide present, no effects are to be expected of these manipulations. The surgery and injections are thus of no significance for the purpose of the current study, for which we were solely interested in behavioral patterns of copulating rats.

Rats were housed in Macrolon IV^®^ cages on a reversed 12h light/dark cycle (lights on between 23:00 and 11:00) in a room with controlled temperature (21 ± 1 °C) and humidity (55 ± 10%), with *ad libitum* access to standard rodent food and tap water. Animals were housed in same-sex pairs with exception of a one-week post-surgery recovery period during which males were single-housed. Males underwent 3 sexual training sessions (once a week) before behavioral testing.

A total of 36 female Wistar rats were ovariectomized as previously described^15^ and used as stimulus animals during the copulation sessions. Briefly, a medial dorsal incision of the skin of about 1 cm was made, and the ovaries were located through a small incision in the muscle layer on each side. The ovaries were extirpated and a silastic capsule containing 10% 17β-estradiol (Sigma, St. Louis, USA) in cholesterol (Sigma, St. Louis, USA) was placed subcutaneously through the same incision. The muscle layer was sutured and the skin was closed with a wound clip. One week of recovery was allowed before the females were used in a copulation session. Four hours before behavioral assessment, female rats were subcutaneously injected with 1 mg progesterone (5 mg/mL; Sigma, St. Louis, USA) in peanut oil (Apotekproduksjon, Oslo, Norway)) to induce receptivity.

#### Groningen

29 male Wistars Unilever (Envigo, Venray, the Netherlands) Rats (approximately 7-8 months old) were housed under reversed 12h light/dark cycle (lights on between 20:00 and 08:00) with *ad libitum* access to food and water. Males underwent behavioral assessment weekly for 7 weeks.

Forty female rats were tubal ligated in order to prevent pregnancies. To perform tubal ligation surgery, females were anesthetized (Isoflurane) and given pain relief (Fynadine, 0.1 mg/100 g) before surgery, and 24 and 48 h after surgery. Females were at least 12 weeks old when surgery was performed, and 2 weeks of recovery were given before receptivity was induced with estradiol (50 μg in 0.1 ml oil, S.C.) 36–48 h before the copulation test. Females were used not more than once in 2 weeks and not more than two times per experimental day.

#### Texas

The data from this cohort comes from two different batches of Long-Evans males (Envigo, Indianapolis, IN, USA); 8 males were approximately 7-8 months old, and 4 males were approximately 3-4 months old during the experiment. Rats were pair housed with same-sex cage mates in hanging polycarbonate cages. The animals were kept on a reversed 12h light/dark cycle (lights on between 22:00 and 10:00) in a room with controlled temperature and humidity, with *ad libitum* access to standard rodent food and tap water. The eight older males in this cohort had previously gained sexual experience as stud males in a female paced-mating set-up. The four younger males were trained in the copulation test set up once per week for three weeks prior to observations for the present study.

Ten Long-Evans females (Envigo, Indianapolis, IN, USA) were ovariectomized at least one week before any behavioral testing took place and used as stimulus animals. To induce sexual receptivity, females were subcutaneously administered 10 μg of estradiol benzoate (Sigma, St. Louis, USA) in sesame oil 48 hours prior to the copulation test, and 1 mg of progesterone (Sigma, St. Louis, USA) in sesame oil 4 hours prior to the copulation test.

The males in the Tromsø, Groningen, and Texas cohorts were selected on the basis of the occurrence of at least one post-ejaculatory interval within a standard 30-minute copulation test.

### 2.2 Copulation test

#### Tromsø

Male subjects were assessed in the copulation test directly after being tested in the sexual incentive motivation test (as part of a previous study^14^). The sexual incentive motivation test consists of a 10-minute free exploration of an arena and socio-sexual stimulus animals that are not accessible for contact interaction. The male subjects were habituated to the sexual incentive motivation test and so no effects on the copulation tests are to be expected. The copulation test, and focus of the current study, was conducted in rectangular boxes (40 × 60 × 40 cm) with a Plexiglas front filled with regular wood chips, in a room with lights on. A receptive female was placed in the copulation box, after which the experimental subject was introduced to start the test.

#### Groningen

The copulation test occurred in wooden rectangular (57 cm × 82 cm × 39 cm; glass wall) boxes with regular wood chips covering the floor, in a room with red light. Rats habituated for 10 min to the testing box right before the test session. After the habituation period, a receptive female was introduced into the box, which started the test.

#### Texas

The copulation test was conducted in rectangular plexiglass boxes (37 × 50 × 32 cm) with regular bedding material (Aspen wood shavings) covering the floor, in a room with red light. A receptive female was placed in the copulation box, after which the experimental subject was introduced and the test was started.

All copulation tests in all labs were conducted during lights-off time, lasted for 30 minutes, and were recorded on camera. Behavior was later assessed from video.

### 2.3 Behavioral assessment

#### Tromsø

Copulation tests were assessed from session 4 (half of the males) and session 5 (half of the males). Males had thus gained sexual experience during 3 or 4 sessions prior to assessment. Behavioral annotation was done for the first ejaculation series (i.e., until the first mount or intromission after the first post-ejaculatory interval).

#### Groningen

Copulation tests were assessed from session 4 (half of the males) and session 5 (half of the males). Males had thus gained sexual experience during 3 or 4 sessions prior to assessment. In addition, session 7 (i.e., after an additional 2-3 sessions of sexual experience allowance) was assessed for all of the males. Behavioral annotation for all of the sessions was done for the first ejaculation series, as well as for the second ejaculation series if 2 post-ejaculatory intervals occurred during the 30-minute test.

#### Texas

Eight of the males in the Texas cohort had previously gained extensive sexual experience as stimulus animals during tests of paced mating behavior. Session 2 of the copulation tests as described was used for assessment of these animals. The remaining four animals only gained sexual experience in the copulation test, and behavioral assessment was done from session 5 (these animals had thus gained sexual experience during 4 sessions prior to assessment). Behavioral annotation for all of the males was done for the first ejaculation series, as well as for the second ejaculation series if 2 post-ejaculatory intervals occurred during the 30-minute test.

#### All cohorts

Behavioral assessment consisted of scoring behavioral events by means of the Observer XT version 12 software (Noldus, Wageningen, the Netherlands). For 1 (Tromsø) or 2 ejaculation series (Groningen and Texas) we behaviorally annotated 100% of the elapsed time according to the following ethogram: the copulatory behaviors mount, intromission, and ejaculation; clasping (mounting the female without pelvic thrusting); genital grooming (grooming of own genital region); other grooming (autogrooming in other regions than genital); chasing (running after the female); anogenital sniffing (sniffing the anogenital region of the female); head towards female (head oriented in the direction of the female while not engaging in other behavior); head not towards female (any behavior that is not oriented towards the female except grooming, such as walking, sniffing the floor, standing still with head direction away from female). For mount bout and time out analysis, the definition as posed by Sachs and Barfield was employed^2^: “A sequence of mounts (one or more), with or without intromission, uninterrupted by any behavior (other than genital autogrooming) that is not oriented towards the female”. Mount bouts and time outs during the copulatory tests were identified through review of the events between copulatory behaviors (mounts, intromissions, and ejaculations). If any other behavior other than genital grooming or “head towards female” occurred between copulatory behaviors, this marked the end of one mount bout (i.e., time of the end of the last copulatory behavior) and beginning of the next mount bout (i.e., time of the next copulatory behavior), and the time in between as a time out duration (see Figure 1A for a schematic overview). From these data points the outcome measures as listed in table 1 were determined (see also ^16^).

**Figure 1.**
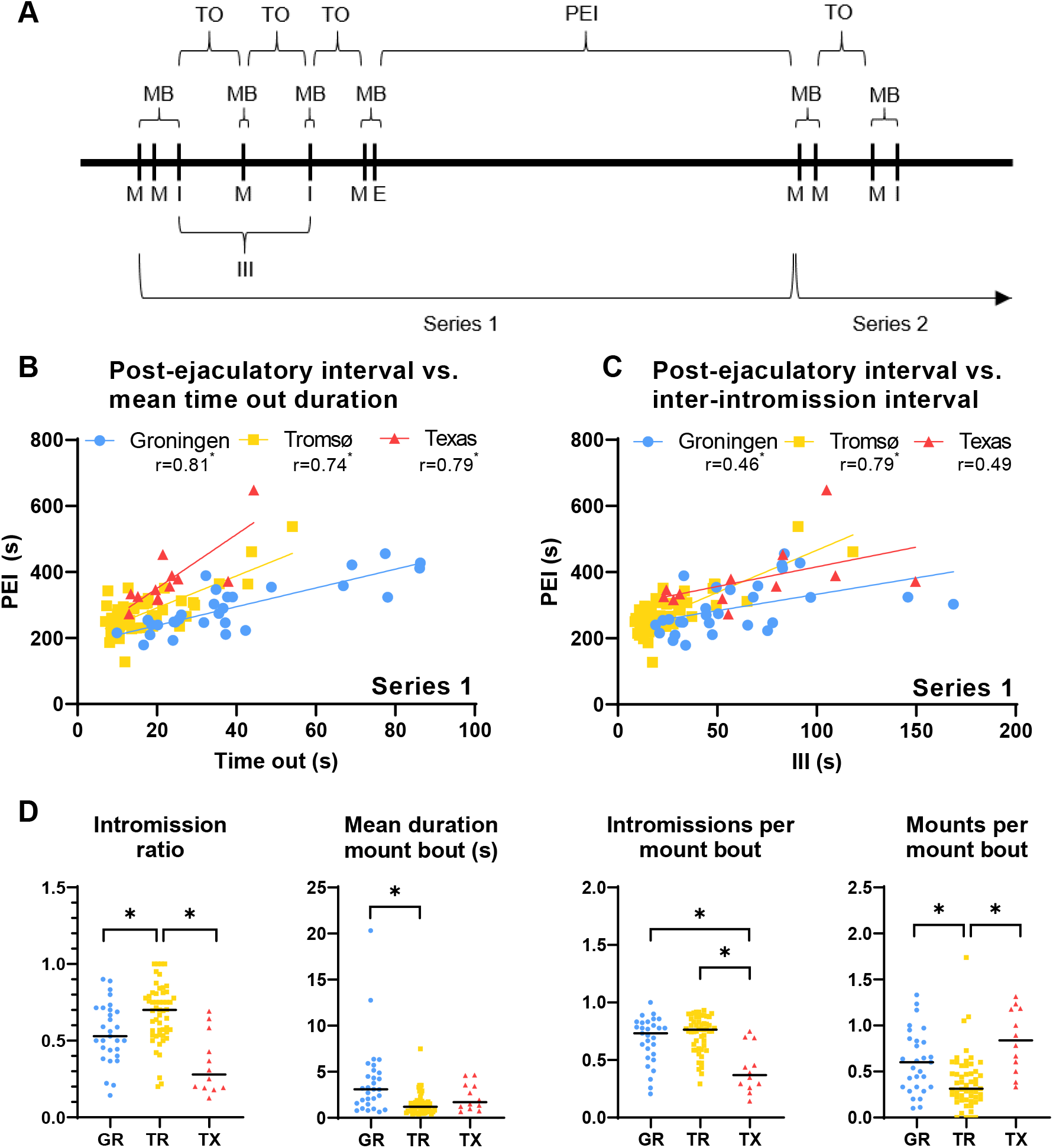
The post-ejaculatory interval correlates with mean time out. **(A)** Schematic overview of male sexual behavior organization. M; mount, I; intromission, MB; mount bout, TO; time out, III; inter-intormission interval, PEI; post-ejaculatory interval. **(B)** Correlation of post-ejaculatory interval and mean time out duration for ejaculation series 1 for Groningen, Tromsø, and Texas cohorts. **(C)** Correlation of post-ejaculatory interval and inter-intromission interval for ejaculation series 1 in Groningen, Tromsø and Texas cohorts. **(D)** Copulation parameters for all cohorts: intromission ratio, mean mount bout duration, mean number of mounts per mount bout, and mean number of intromissions per mount bout. Horizontal lines; median. **All panels:** n=29; 53; 12, *p<0.05.

**Table 1.** Copulation test outcome measure definitions

| Outcome measure | Definition |
| --- | --- |
| Latency to first mount or intromission | Time from the start of the test to the first mount or intromission |
| Number of mounts | Total number of mounts preceding ejaculation |
| Number of intromissions | Total number of intromissions preceding ejaculation |
| Intromission ratio | Number of intromissions in the ejaculation series divided by the total number of copulatory behaviors (mounts + intromissions) in the ejaculation series |
| Number of mount bouts | Total number of mount bouts preceding ejaculation |
| Mounts per mount bout | Mean number of mounts per mount bout in an ejaculation series |
| Intromissions per mount bout | Mean number of intromissions per mount bout in an ejaculation series |
| Mount bout duration | Time from the first copulatory behavior in a mount bout until the first behavior within the following time out |
| Time out duration | Time from the end of one mount bout to the start of the next mount bout |
| Inter-intromission interval | Time between intromissions in an ejaculation series, calculated from the first intromission |
| Latency to ejaculation | Time from the first mount or intromission to ejaculation |
| Post-ejaculatory interval | Time from the first ejaculation to the next copulatory behavior (mount or intromission) |

### 2.4 Data analysis and statistics

#### Correlation between PEI, III and time outs

The post-ejaculatory interval versus mean time out duration and the inter-intromission interval for the corresponding ejaculation series were analyzed with Pearson correlation coefficients. The mean time out duration was calculated for each subject from all time outs in the corresponding ejaculation series.

#### Analysis of copulation and mount bout characteristics

The behavioral data used for comparisons between cohorts were not normally distributed and were therefore analyzed with non-parametric tests. The Kruskal-Wallis test followed by Dunn’s multiple comparison posthoc test was employed for comparisons between copulation test outcome parameters of the three different cohorts.

#### Within-subject consistency within and across copulatory sessions

The Wilcoxon matched-pairs signed rank test was used to analyze the data for ejaculation series 1 compared to ejaculation series 2 in the Groningen and Texas cohorts. Pearson correlation coefficients was employed to analyze the relation of PEI and time out in the different ejaculation series, as well as to analyze the relation of PEI/time out in ejaculation series 1 and PEI/time out in ejaculation series 2.

#### Time out predictors

The duration of each mount bout versus the duration of its following time out was analyzed with Pearson correlation coefficients. For comparison of data corresponding to individual mount bouts/time outs, data points were z-scored within each rat using the following calculation: z-score = ((data point) – (mean of the data points for the rat))/(standard deviation of the data points for the rat). Z-scores of the different cohorts were then analyzed by means of Mann Whitney U tests in case of 2 groups, or Kruskal-Wallis and Dunn’s posthoc tests for 3 or more groups. For the time-binned analysis of time out duration, the first 33% of time outs were defined as time-bin 1, the second 33% as time-bin 2, and the last 33% as time-bin 3. For the time out duration per mount bout stimulation analysis, mount bout types with less than 10 data points were excluded from analysis (e.g. 3 mounts, 2 intromissions).

The behavioral data were extracted from the Observer data files and analyzed using custom Python 3.8 scripts. The scripts are available for sharing upon request. All statistical analyses were performed in GraphPad Prism version 9.0.0 (GraphPad Software, San Diego, CA, USA). In all cases, alpha was set at 0.05 and tests were two-tailed.

## 3. Results

### 3.1 Relation of inter-copulatory intervals

#### Correlation between PEI, III and time outs

Our analysis first focused on how inter-copulatory intervals, i.e. the post-ejaculatory interval (PEI), time outs, and inter-intromission interval (III), relate to one another. Our mount bout-based analysis (Fig. 1A) showed that the PEI was strongly correlated with the mean time out duration in all of the cohorts: Groningen (Figure 1 B; *r*=0.81, p<0.001), Tromsø (Fig. 1B; *r*=0.74, p<0.001), and Texas (Fig. 1B; *r*=0.79, p=0.002). Correlation between the PEI and the III was also strong in the Tromsø cohort (Fig. 1C; *r*=0.79, p<0.001), but weak in the Groningen cohort (Fig. 1C; r=0.46, p=0.01), and not significant in the Texas cohort (Fig. 1C; r=0.49, NS).

#### Analysis of copulation and mount bout characteristics

We next examined whether the difference in correlation strength of PEI vs. III between the cohorts could be explained from copulatory parameters. All copulatory parameters and comparisons between the three cohorts can be found in Supplementary Table 1. Only those parameters that are relevant for the current assessment will be discussed in this section. We hypothesized that PEI vs. III correlation is stronger in cohorts in which the III resembles the mean time out duration. If each mount bout consists of only a single intromission, mean time out duration and III are the same. Thus, III more strongly approaches mean time out duration in cohorts with a high number of intromissions, short mount bouts, and more mount bouts with an intromission and relatively few mounts. We found that the copulatory parameters intromission ratio (i.e., number of intromissions divided by total number of copulatory behaviors) (Fig 1D; *H(2)*=21.67, p<0.001), mean duration of mount bout (Fig 1D; *H(2)*=20.30, p<0.001), mean number of mounts per mount bout (Fig 1D; *H(2)*=20.47, p<0.001), and mean number of intromissions per mount bout (Fig 1D; *H(2)*=17.51, p<0.001)in the first ejaculation series differed significantly between the cohorts. The Tromsø cohort had a larger intromission ratio (Fig 1D; p<0.001), more intromissions per mount bout (Fig 1D; p<0.001), and less mounts per mount bout (Fig 1D; p<0.001) than the Texas cohort. The Tromsø cohort also had a larger intromission ratio (Fig. 1D; p=0.023), a shorter mean mount bout duration (Fig 1D; p<0.001) and less mounts per mount bout (Fig. 1D; p=0.007) than the Groningen cohort. The Groningen cohort had more intromissions per mount bout than the Texas cohort (Fig 1D; p=0.003). These results show that correlation between PEI and III is indeed stronger when time out and III are similar, as is the case in the Tromsø cohort, and explains why the PEI and III correlated stronger in this cohort than in the other cohorts.

#### Within-subject consistency within a copulatory session

To see whether the mean PEI duration and mean time out duration followed the same pattern over time within the same rats, we looked at how these parameters change from the first ejaculation series to the second ejaculation series within a copulation session, and over different copulation sessions. In the Groningen cohort, the PEI (Fig. 2A; *W*=−251, p<0.001) as well as the mean time out duration (Fig. 2B; *W*=−129, p=0.036) increased in the second ejaculation series compared to the first ejaculation series. We did not find statistically significant effects in the Texas cohort for ejaculation series 1 compared to ejaculation series 2. The PEI of ejaculation series 2 also correlated with the mean time out duration in ejaculation series 2 in the Groningen cohort (Fig. 2C; *r*=0.70, p=0.002) as it did in ejaculation series 1. There was moderate correlation of PEI and mean time out duration in ejaculation series 2 in the Texas cohort, but this was not statistically significant (Fig. 2C; *r*=0.66, NS).

**Figure 2.**
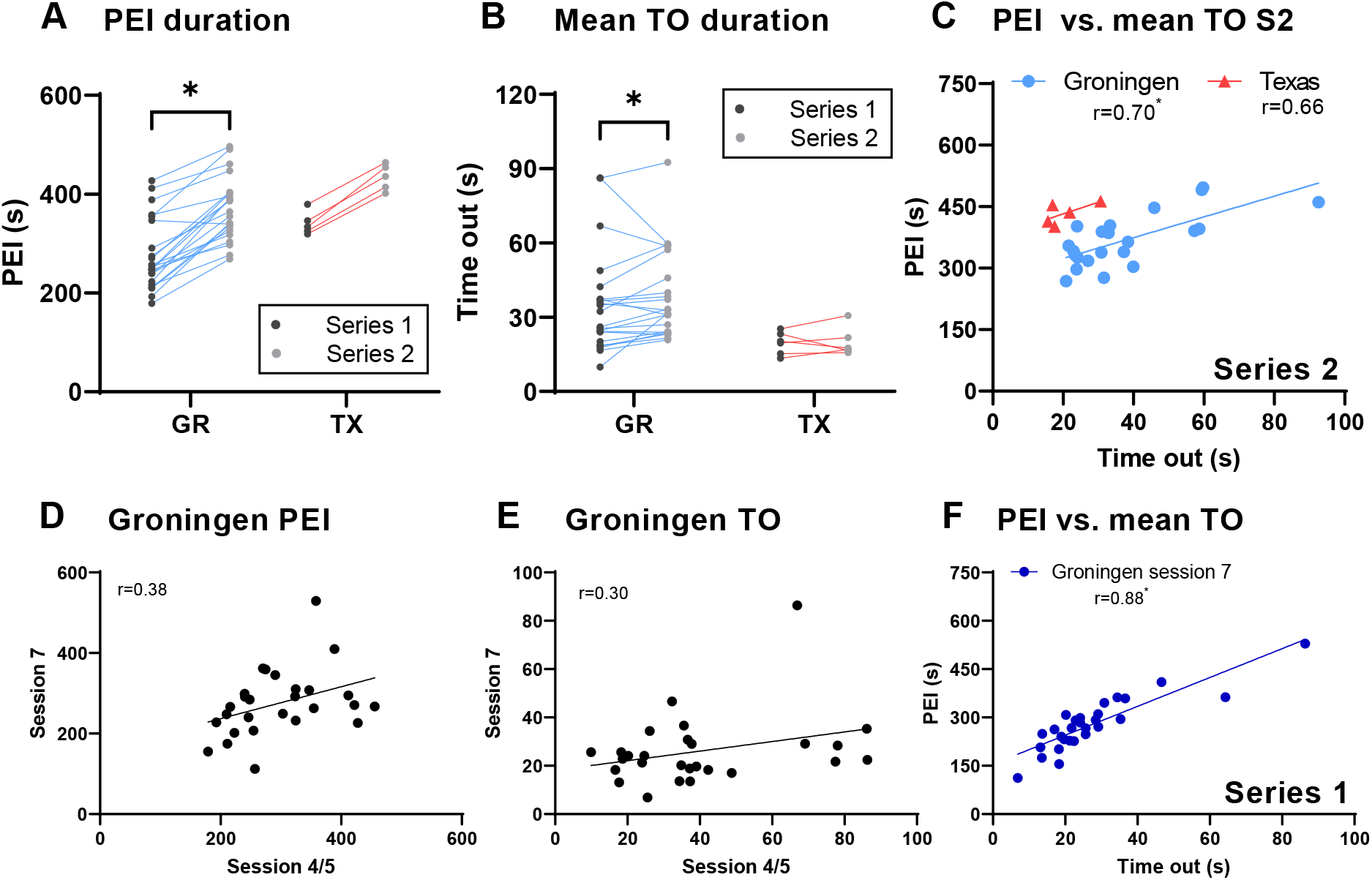
Post-ejaculatory interval and time out both increase over ejaculation series. **(A)** Post-ejaculatory interval duration in ejaculation series 1 compared to ejaculation series 2 within the same animals from the Groningen and Texas cohorts, n=22; 5. **(B)** Mean time out duration in ejaculation series 1 compared to ejaculation series 2 within the same animals from the Groningen and Texas cohorts, n=22; 5. **(C)** Correlation of post-ejaculatory interval and mean time out duration for ejaculation series 2 in the Groningen and Texas cohorts, n=22; 5. **(D)** Correlation of post-ejaculatory interval in copulation session 4/5 with copulation session 7 within the same Groningen animals, n=27. **(E)** Correlation of mean time out duration in copulation session 4/5 with copulation session 7 within the same Groningen animals, n=27. **All panels:** PEI; post-ejaculatory interval, TO; time out, *p<0.05

#### Within-subject consistency across copulatory sessions

Both the PEI and the mean time out duration in the first ejaculation series did not show significant correlation from one copulation session (the 4^th^ or 5^th^ occasion of copulation) to another copulation session (the 7^th^ occasion of copulation) in the Groningen cohort (Fig. 2D-E). However, the correlation of PEI and mean time out duration in the first ejaculation series was persistent over multiple copulation sessions, as the effect was still present and of the same magnitude in the later copulation session of the Groningen cohort (Fig. 2F; r=0.88, p<0.001). Thus, PEI and mean time out duration vary over copulation sessions within rats, but the correlation of the two parameters within each copulation session is consistent.

### 3.2 Time out predictors

We next assessed whether any mount bout characteristic predicted the duration of the subsequent time out. First, the duration of individual mount bouts did not correlate with the duration of the subsequent time out (Fig. 3A). Second, we considered that copulatory pace might be faster or slower depending on how close the male is to ejaculation. Therefore, we examined whether individual time out duration is dependent on the relative time point within the ejaculation series. We divided the ejaculation series into three-time bins, each consisting of a third of the total number of time outs within the ejaculation series, and analyzed whether standardized (z-scored within subject) time out duration differs between time bins for each of the cohorts. We found that standardized time out duration was different over time bins in the Groningen cohort (Fig. 3B; *H(2)*=28.28, p<0.001): the median time out duration was longer in the third time bin compared to both the second (Fig. 3B; p<0.001) and the first time bin (Fig. 3B; p<0.001). We did not find this effect in the Tromsø or Texas cohort (Fig. 3B).

**Figure 3.**
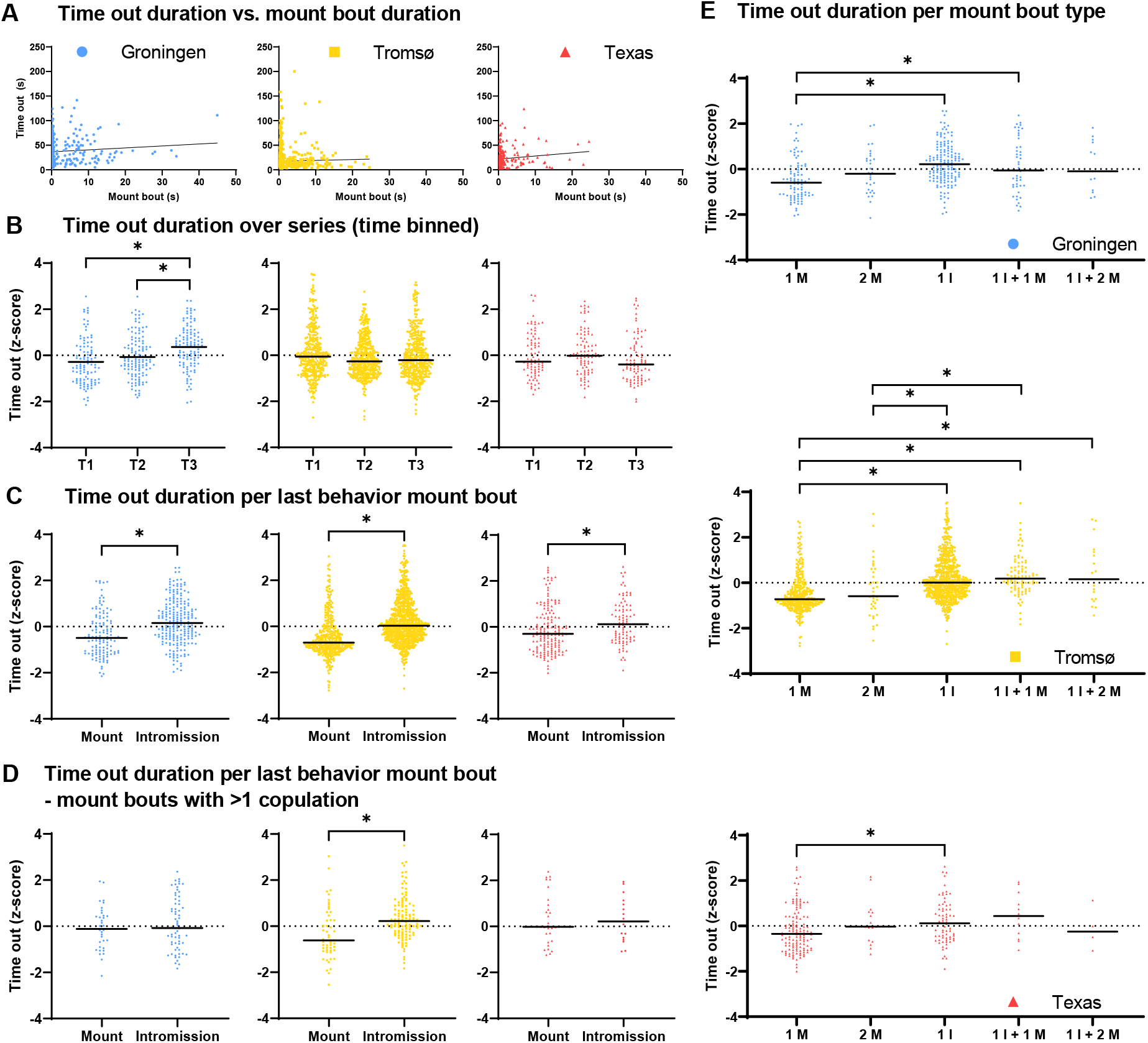
More stimulation within mount bout is associated with longer time out. **(A)** Correlation of individual mount bout duration with subsequent time out duration in Groningen, Tromsø and Texas cohorts, n=341; 1118; 249. **(B)** Z-scores of individual time out durations during the first, second, and third third of the ejaculation series; Groningen (n=103; 123; 114), Tromsø (n=354; 388; 376) and Texas (n=79; 84; 83). **(C)** Z-scores of individual time out duration after mount bouts with mount vs. intromission as last copulation; Groningen (n=124; 216), Tromsø (n=366; 751) and Texas (n=160; 86). **(D)** Z-scores of individual time out duration after mount bouts consisting of multiple copulations with mount vs. intromission as last copulation; Groningen (n=35; 64), Tromsø (n=52; 111) and Texas (n=28; 16). **(E)** Z-scores of individual time out durations after mount bouts with different total copulatory stimulation; Groningen (n=88; 30; 152; 45; 13), Tromsø (n=314; 36; 640; 87; 18) and Texas (n=132; 16; 70; 12; 4), M; mount, I; intromission. **All panels:** Horizontal lines; median, *p<0.05.

Third, we assessed whether mount bouts that end in an intromission might induce a longer time out than mount bouts that end in a mount. We found that the median duration of time outs that follow a mount bout ending with an intromission was shorter than the duration of time outs that follow a mount bout ending with a mount in the Groningen (Fig. 3C; *U*=8834, p<0.001), Tromsø (Fig. 3C; *U*=72940, p<0.001), and Texas (Fig. 3C; *U*=5496, p=0.009) cohorts. To examine whether this effect of the last behavior within a mount bout is independent of the number of copulations within the mount bout, we ran the same analysis after exclusion of all time outs that followed mount bouts consisting of only a single copulatory behavior. This analysis showed that the effect disappeared in the Groningen and Texas cohorts, but remained in the Tromsø cohort (Fig. 3D; *U*=1739, p<0.001). We also noted that of the mount bouts with multiple copulations, only 5 out of 117 (4.3%) mount bouts that ended in a mount also contained an intromission (data not shown). This is consistent with our observation that 1057 out of 1068 intromissions (99%) in the full data set ended the mount bout (data not shown). These results indicated that the significant effects of the last behavior within a mount bout on the subsequent time out duration might be a function of the sum of components of the mount bout, which might rather be the true predictor of time out duration.

Fourth, based on the conclusion above, we hypothesized that the total sensory stimulation within the mount bout predicts the following time out duration. We defined mount bout types by the sum of copulatory components of the mount bout and compared the duration of following time outs between mount bout types (bouts with a) 1 mount, b) 2 mounts c) 1 intromission, d) 1 intromission + 1 mount, e) 1 intromission + 2 mounts. Indeed, there was a significant difference between the median standardized time out duration after different mount bout types in all of the cohorts: Groningen (Fig. 3E; *H(4)*=34.30, p<0.001), Tromsø (Fig. 3E; *H(4)*=162.0, p<0.001), and Texas (Fig. 3E; *H*(4)=11.6, p=0.020). Time outs following mount bouts of 1 mount had a shorter median duration than time outs following mount bouts of 1 intromission in the Groningen (Fig. 3E; p<0.001), Tromsø (Fig. 3E; p<0.001), and Texas (Fig. 3E; p=0.046) cohorts. This was also the case for 1 mount compared to 1 intromission and 1 mount in the Groningen (Fig. 3E; p=0.013) and the Tromsø cohort (Fig. 3E; p<0.001), and compared to 1 intromission and 2 mounts in the Tromsø cohort (Fig. 3E; p=0.009). In addition, time outs following a mount bout of 2 mounts had a shorter mean duration compared to time outs following mount bouts of 1 intromission (Fig. 3E; p=0.008) and compared to 1 intromission and 1 mount (Fig. 3E; p=0.001) in the Tromsø cohort only. Additional mounts in mount bouts with intromissions did not lengthen the subsequent time out any further in any of the data sets (Fig. 3E).

## 4. Discussion

In behavioral neuroscience, it is important to know as much about the structure and organization of the behavior under investigation as possible, because a detailed understanding of the behavior lends itself to better assessment of causal neurobiological mechanisms underlying the behavior. In light of this, the advancement of research on male rat sexual behavior has been disappointing in the recent decades, as behavioral assessment of copulation is most often reduced to the annotation of mounts, intromissions, and (usually one) ejaculation only. Unfortunately, the pioneering study by Sachs and Barfield on temporal patterning of male rat copulation has not had a lasting impact. The relationship between, and predictors of inter-copulatory intervals in male rat sexual behavior have yet to be elucidated. In the current study, we showed that the PEI is strongly correlated with mean time out duration and that time out duration is at least partially predicted by the total sensory stimulation in the preceding mount bout. These conclusions are remarkable because they were observed in three different cohorts of rats, of two different strains, and in rats of different ages from different origins. In addition, the experiments were carried out in three different laboratories in different geographical locations, with slightly different procedures. This emphasizes the generalizability of our results in the context of male-paced copulation in rats. Our findings advance our understanding of how the PEI and time out are related and possibly regulated by a similar central neuronal inhibition. Our results show how a more detailed analysis of behavioral structure and organization can provide valuable insights for future research.

### 4.1 Relation of inter-copulatory intervals

That the PEI and the III, the most common measure of temporal patterning in recent literature, are related was already apparent from factor analyses in which these parameters load onto the same factor.^5,6^ However, as mount bouts and time outs are a better measure of the natural temporal patterning than III in male rat copulation,^2^ our finding that time outs have a stronger correlation with PEI than III is logical. The III disregards mounts even though they are central copulatory behaviors and contribute to the facilitation of ejaculation,^11^ and is strongly dependent on the intromission ratio, or efficiency, of the male. Still, in our data set, PEI and III are also strongly correlated in the Tromsø cohort. This can be explained by the notion that this cohort had a high intromission ratio, a short mount bout duration, relatively few mounts per mount bout, and at least 1 intromission in the majority of mount bouts. These copulation characteristics make for the mean time out duration to strongly approximate the III, which would be much larger than the time out when more mounts and less intromissions occur per mount bout. This explains the strong PEI and III correlation in the Tromsø cohort and emphasizes how this correlation is dependent on how closely mean time out duration resembles III. We stress again that mount bout-based analysis should be standard for assessment of copulatory pace of male rat copulation and that III is not sufficient for this goal. As an example for the general utility of mount bout-based analysis, we were recently able to draw more informed conclusions about the cause of increased ejaculation latency upon a manipulation.^14^ Because there was no effect on time out duration, and thus on copulatory pace, in this study, we could state that the prolonged ejaculation latency was caused by a decreased sensitivity to reach ejaculation threshold. In order to advance the field of sexual behavior further, it is vital to have a better behavioral understanding in depth and to measure the parameters of the natural organization of copulation in the form of mount bouts and time outs.

It has previously been shown that the PEI increases over each following ejaculation series when males copulate to exhaustion,^10^ whereas the III follows a U-shape, and no data to our knowledge has been published on time out. Since PEI and time out strongly correlates with mean time out duration, and not reliably with III, it would follow logically that mean time out would follow a similar pattern over ejaculation series as the PEI. We indeed found that the PEI and mean time out duration in our Groningen cohort was longer in the second ejaculation series than in the first. There was a similar trend in the Texas cohort, but this was not statistically significant due to a much smaller sample size. We did not have the data to investigate the course of the time out over more than two ejaculation series, but it would be interesting if future research could focus on analysis of males copulating to exhaustion, yielding more ejaculation series to study trends over time. The strong correlation between PEI and mean time out, together with the fact that both of these parameters increase over ejaculation series, suggests that the orchestration of both these intervals on the neurobiological level could be related.

### 4.2 Within-subject consistency of inter-copulatory intervals

We found that both the PEI and the time out are not correlated across copulation sessions within the same cohort of males. Thus, temporal patterning of copulation, as measured by inter-copulatory intervals, varies from session to session. This is consistent with the fact that the PEI does not seem to be a part of sexual behavior endophenotypes in male rats, as rapid ejaculators do not have a shorter PEI than normal ejaculators.^17^ Importantly, even though the PEI and time out vary over copulation sessions, the strong correlation between these two parameters was consistent over sessions, indicating that their variation is unidirectional over sessions within a rat. All of the males in our cohorts were allowed sufficient recovery after each copulation session as they were behaviorally tested only once per week, which is enough for all copulation parameters to return to baseline even after exhaustion.^18^ We hypothesize that variability over sessions in copulatory pace (as determined by PEI and time out) could simply be caused by daily condition of the male, or is perhaps dependent on the female stimulus. In all of our labs, females are paired with males at random and replaced in case of signs of reduced receptivity or sexual rejection (either of which rarely occur). It has been shown that the III and number of intromissions in the first ejaculation series (as well as a trend for PEI) in a semi-natural environment are different when domesticated males mate with females of the same strain versus females that are caught in the wild.^19^ In the same study it was demonstrated that 84% of female paracopulatory behavior episodes are followed by an intromission, whereas only 13% of male-initiated copulations (i.e. not preceded by female paracopulatory behavior) resulted in an intromission. The authors note that differences in paracopulatory behavior frequency seems to account for the copulation difference of males mating with the same strain versus with a wild female.^19^ It needs to be addressed though, that males and females initiate copulation in a semi-natural environment just as often, and that the occurrence of copulatory acts is not mainly controlled by the female.^20^ Even though copulatory pacing is thus shared between males and females in a semi-natural environment and seemingly controlled by the male in the standard copulation apparatus, these findings indicate that females are capable of exerting some control over male copulation speed and efficiency. Additional evidence for this notion comes from an experiment in which females were removed after each mount bout and returned to the copulation apparatus upon a bar press.^21^ Under these circumstances, the mean time out duration increased, suggesting a stimulatory role of the presence of the female on the reinitiation of mounting by the male. This effect might still be minimal in an *ad libitum* male-paced setting, but possibly relevant for slight session-to-session variability in inter-copulatory intervals if there are individual differences between stimulatory properties of the female in the context of male-paced mating, perhaps found in the number of paracopulatory behaviors displayed by the female. It would be interesting to further investigate the role of the female in male-paced mating protocols.

### 4.3 Mount bout predictors of following time out duration

Because the III first increases and then decreases over time within the ejaculation series,^22^ we examined how the time out duration is distributed within the first ejaculation series. We found that time outs in the third time bin of the ejaculation series in the Groningen cohort were significantly longer than in the first- and second- time bins. We did not find this effect in the other cohorts. A possible explanation for this discrepancy could be that in Groningen, the male subject has a 10-minute habituation to the copulation box (not cleaned) before the female is introduced, whereas in Tromsø and Texas the male is introduced after the female. The habituation in the copulation box, soiled with pheromones and odors, could have increased sexual arousal before introduction of the female, leading to a shortening of time outs in the start of the copulation test and a gradual normalization over the ejaculation series. This is in line with the fact that we did not find effects of time bin on time out duration in the second ejaculation series of the Groningen cohort (Suppl. Fig. 3A). Overall, the time out does not consistently vary over time within the ejaculation series as the III does, but more research into the role of sexual arousal on time out would be interesting

Next, we showed that time outs following mount bouts that ended with an intromission were longer than time outs following mount bouts that ended with a mount. Pollak and Sachs (1976) have reported a similar assessment from two cohorts of males (n=7 and n=5) and found that time out duration after mount bouts that ended with an intromission was increased by 26% and 9% for the two replicates respectively, although not statistically significant.^13^ Our data shows a similar magnitude of time out duration increase, but we did find a statistically significant effect in our data set. Because we analyzed on the level of individual time out that was standardized for each rat by z-scoring, instead of analyzing the average for each subject rat, our data set consists of a much larger sample size. The advantage of this approach is that the z-score better reflects the difference in duration between time outs within a rat, while making it possible to still analyze on a group level and compare between different cohorts. This difference in approach compared to Pollak and Sachs, and our much larger number of male subjects, could account for our different statistical outcomes. One other reason that our results reached statistical significance, but not the results from Pollak and Sachs, may be that our data set consisted of a relatively high percentage of mount bouts consisting of only a single copulatory behavior, whereas Pollak and Sachs report a mean number of 1.5 mounts per mount bout in their cohort. This difference in behavioral phenotype may possibly be due to changes in genetic make-up of animals over time. When excluding the mount bouts with a single copulation from analysis, we found a smaller effect of the last event in a mount bout on the following time out. Still, since 99% of intromissions end a mount bout (similar to 90% reported by Pollak and Sachs), mount bouts of multiple copulations ending in a mount are far less likely to include an intromission as well. Therefore, we proceeded with analyzing whether the total stimulation within the mount bout might be the determining factor for the duration of the subsequent time out.

We found that mount bouts consisting of 1 intromission (or 1 intromission and 1 mount in two of the cohorts) induced a longer time out than mount bouts consisting of 1 mount in all of the cohorts. This seems incongruent with earlier reports that show that time out duration distribution is not affected by the prohibition of intromissions.^2,12^ However, males that could not intromit tend to have more mounts per mount bout: from 1.5 to 2.6 upon penile lidocaine application^13^ and from 2.5 to 3.1 and 9.9 when mating with a female with closed vagina or upon penile tetracaine application, respectively, although not statistically significant.^2^ Therefore, if time out duration following mount bouts of 3, 4 or even more mounts is similar to time out duration following mount bouts with at least 1 intromission, it is very possible that no effect would be found of intromission prevention on time out duration distribution. In a data set in which we compiled the data of all three cohorts, we did not find a difference in time out duration following mount bouts of 1 intromission versus mount bouts of three or more mounts (no intromissions), although this data set was small (Suppl. Fig. 4). This underscores that our results are not necessarily in disagreement with the earlier reports and we conclude that time out duration is at least partially under the control of the total stimulation within the preceding mount bout, with a ceiling effect for intromissions.

### 4.4 Reflections on a hypothesis for a shared central mechanism of inter-copulatory intervals

We showed that PEI strongly correlates with time out, that both of these parameters increase in the first ejaculation series compared to the second ejaculation series, and that even though both of these parameters vary across copulation sessions, their correlation remains present in each copulation session. Moreover, time out is longer after mount bouts with more penile sensory stimulation, whereas the longest inter-copulatory interval (i.e, PEI) follows the strongest sensory stimulation (i.e., ejaculation). Our interpretation of these findings suggests a possibility of PEI and time out being under a similar inhibitory neuronal control. However, alternative hypotheses may be considered. First, one possible cause of increased time out duration after mount bouts that contain intromissions could be that intromissions may induce increased duration of genital grooming. While it is indeed true that mean genital grooming duration is longer after intromissions than after mounts, it was also reported in the same previous study that this effect disappears when only mounts that end a mount bout are considered.^23^ Thus, duration of genital grooming after the last behavior within a mount bout is independent of whether that last behavior was an intromission or a mount. In line with this, desensitization of the penis by means of topical application of anesthetic ointment or surgical transection of the penile nerve does not affect genital grooming duration after mounts and intromissions that end a mount bout, suggesting that genital grooming duration is not dependent on the magnitude of sensory feedback within the mount bout.^24^ In addition, prevention of genital grooming does not affect ejaculation latency, mounting and intromission frequency, and PEI duration.^25^ It is hence postulated that genital grooming might rather be part of a motor program of copulatory behavior.^24^ Second, an alternative explanation for the strong correlation of PEI and time out may be that males with a longer PEI simply have more intromissions preceding ejaculation, since mount bouts that contain intromissions induce a longer time out than mount bouts without intromissions. In an extra analysis, we found no correlation whatsoever between PEI duration and the number of intromissions in the first ejaculation series in any of the cohorts (Suppl. Fig. 5). Concluding, our working hypothesis remains that both the PEI and time out are the result of a copulatory inhibition, which is induced by the sum of sensory penile stimulation.

A question that logically arises considering this working hypothesis is whether there is a refractory period after a mount bout like after an ejaculation. The PEI is known to consist of two phases: the absolute refractory period (the first 75% of the PEI duration) and the relative refractory period (the last 25% of the PEI duration). During the relative refractory period, males can be moved to reinitiate copulation faster through non-specific stimulations that presumably increase general arousal, such as handling,^26^ electrical shock,^27,28^ and removal of the female for short periods of time.^9^ These interventions have no effect during the absolute refractory period. If PEI and time out share common mechanisms, one might expect that the time out also consists of an absolute and relative refractory period. There is some evidence for this. Like the PEI, the time to next copulation after an intromission can be decreased by shortly removing the female,^9^ handling,^26^ or by applying electrical shock after an intromission (as described in Sachs and Barfield (1976)^29^). Interestingly, whereas male rats that have a natural fast copulatory pace (III of less than 30 seconds) are responsive to shocks within 3, 6, 12, or 24 seconds after intromission, naturally slower subjects are unresponsive to shocks within 3 seconds and only marginally responsive to shocks within 6 seconds (as described in Sachs and Barfield (1976)^29^). This is perhaps an indication of an absolute refractory period during inter-copulatory intervals that occur before ejaculation. Future research might provide insight into whether an absolute refraction indeed exists and whether it can be identified as a certain time percentage of a time out.

Another clue about the mechanistic relationship between inter-copulatory intervals and PEI is found in an electrophysiological study. Kurtz and Adler showed that all ejaculations and almost all intromissions are followed by a decrease in hippocampal theta frequency and a desynchronization of hippocampal activity.^30^ Mounts, on the other hand, are followed by a theta frequency decrease in 27% of the cases, but by a theta frequency increase in 73% of the cases. Since intromissions almost always end a mount bout and the chance for a mount to end a mount bout is much smaller, it could be hypothesized that the slowing and desynchronization of hippocampal activity might be at the basis of copulatory inhibition, while increased hippocampal theta frequency is indicative of a continuation of copulation (i.e. the mount bout). Studying these oscillations in the context of a mount bout analysis should answer whether the theta frequency increase indeed only happens after the last behavior in a mount bout, and not after copulations within a mount bout, as well as whether similar electrophysiological patterns can be observed throughout the PEI and the time out. Future research should aim to determine whether inter-copulatory intervals indeed share a central mechanism.

### 4.5 Conclusion

We conclude that PEI and mean time out duration are strongly correlated, and that the total stimulation within a mount bout predicts the length of the following time out. These results were consistent over three different cohorts, despite differences in strain, age, lab, and testing procedure. We hypothesize that both PEI and time out could be regulated by a similar central copulatory inhibition that is at least partially under the control of the magnitude of sensory stimulation. Future research should aim to elucidate the underlying inhibitory mechanisms of both PEI and time out. Moreover, we advocate that the assessment of sexual behavior in male rats should be more extensive and include analysis based on mount bouts, in order to understand measured effects on a more detailed level.

## Acknowledgements

Financial support was received from Norwegian Research Council; grant #251320 to EMS. We thank Josien A. Janssen for data gathering. We thank Carina Sørensen, Katrine Harjo, Ragnhild Osnes, Remi Osnes and Nina Løvhaug for the excellent care of the animals.

## Data sharing statement

The data that support the findings of this study are available from the corresponding author upon reasonable request.

## References

1. Krakauer JW, Ghazanfar AA, Gomez-Marin A, MacIver MA, Poeppel D. Neuroscience Needs Behavior: Correcting a Reductionist Bias. Neuron. 2017;93(3):480–490.

2. Sachs BD, Barfield RJ. Temporal Patterning of Sexual Behavior in Male Rat. J Comp Physiol Psych. 1970;73(3):359–364.

3. Sachs BD, Garinello LD. Interaction between Penile Reflexes and Copulation in Male Rats. J Comp Physiol Psych. 1978;92(4):759–767.

4. Seizert CA. The neurobiology of the male sexual refractory period. Neurosci Biobehav Rev. 2018;92:350–377.

5. Pfaus JG, Mendelson SD, Phillips AG. A correlational and factor analysis of anticipatory and consummatory measures of sexual behavior in the male rat. Psychoneuroendocrinology. 1990;15(5-6):329–340.

6. Dewsbury DA. Factor analyses of measures of copulatory behavior in three species of muroid rodents. J Comp Physiol Psych. 1979;93(5):868–878.

7. Beach FA. Characteristics of masculine “sex drive”. In: Jones MR, ed. Nebraska symposium on motivation. Lincoln: University of Nebraska. 1956.

8. Flannelly KJ, Blanchard RJ, Layng MP, Blanchard DC. Life-span changes in the copulatory behavior of male rats (Rattus norvegicus). J Comp Psychol. 1985;99(1):87–92.

9. Bermant G. Effects of single and multiple enforced intercopulatory intervals on sexual behavior of male rats. J Comp Physiol Psych. 1964;57(3):398–403.

10. Karen LM, Barfield RJ. Differential rates of exhaustion and recovery of several parameters of male rat sexual behavior. J Comp Physiol Psychol. 1975;88(2):693–703.

11. Hard E, Larsson K. Effects of mounts without intromission upon sexual behaviour in male rats. Anim Behav. 1968;16(4):538–540.

12. Lodder J, Zeilmaker GH. Effects of pelvic nerve and pudendal nerve transection on mating behavior in the male rat. Physiology & Behavior. 1976;16(6):745–751.

13. Pollak EI, Sachs BD. Penile movements and the sensory control of copulation in the rat. Behavioral Biology. 1976;17(2):177–186.

14. Huijgens PT, Heijkoop R, Snoeren EMS. Silencing and stimulating the medial amygdala impairs ejaculation but not sexual incentive motivation in male rats. Behav Brain Res. 2021;405:113206.

15. Ågmo A. Male rat sexual behavior. Brain Res Brain Res Protoc. 1997;1(2):203–209.

16. Heijkoop R, Huijgens PT, Snoeren EMS. Assessment of sexual behavior in rats: The potentials and pitfalls. Behavioural Brain Research. 2018;352:70–80.

17. Pattij T, de Jong TR, Uitterdijk A, et al. Individual differences in male rat ejaculatory behaviour: searching for models to study ejaculation disorders. Eur J Neurosci. 2005;22(3):724–734.

18. Jackson SB, Dewsbury DA. Recovery from sexual satiety in male rats. Anim Learn Behav. 1979;7(1):119–124.

19. McClintock MK, Adler NT. The role of the female during copulation in wild and domestic Norway Rats (Rattus norvegicus). Behaviour. 1978;67(1):67–96.

20. Bergheim D, Chu X, Ågmo A. The function and meaning of female rat paracopulatory (proceptive) behaviors. Behav Processes. 2015;118:34–41.

21. Sachs BD, Macaione R, Fegy L. Pacing of copulatory behavior in the male rat: effects of receptive females and intermittent shocks. J Comp Physiol Psych. 1974;87(2):326–331.

22. Dewsbury DA. Changes in inter-intromission interval during uninterrupted copulation in rats. Psychon Sci. 1967;7(5):177–178.

23. Sachs BD, Clark JT, Molloy AG, Bitran D, Holmes GM. Relation of autogrooming to sexual behavior in male rats. Physiology & Behavior. 1988;43(5):637–643.

24. Bitran D, Sachs BD. Penile desensitization does not affect postcopulatory genital autogrooming in rats: evidence for central motor patterning. Physiol Behav. 1989;45(5):1001–1006.

25. Hart BL, Haugen CM. Prevention of genital grooming in mating behaviour of male rats (Rattus norvegicus). Animal Behaviour. 1971;19(2):230–232.

26. Larsson K. Non-specific stimulation and sexual behaviour in the male rat. Behaviour. 1963;20(1):110–114.

27. Barfield RJ, Geyer LA. The ultrasonic postejaculatory vocalization and the postejaculatory refractory period of the male rat. J Comp Physiol Psychol. 1975;88(2):723–734.

28. Caggiula AR, Vlahoulis M. Modifications in the copulatory performance of male rats produced by repeated peripheral shock. Behav Biol. 1974;11(2):269–274.

29. Sachs BD, Barfield RJ. Functional Analysis of Masculine Copulatory Behavior in the Rat. In: Rosenblatt JS, Hinde RA, Shaw E, Beer C, eds. Advances in the Study of Behavior. Vol 7. Academic Press; 1976:91–154.

30. Kurtz RG, Adler NT. Electrophysiological correlates of copulatory behavior in the male rat: evidence for a sexual inhibitory process. J Comp Physiol Psychol. 1973;84(2):225–239.

