## Supplemental data for "Male rat sexual behavior: insights from inter-copulatory intervals"

**Supplementary Table 1** Copulation test outcome parameters for ejaculation series 1 of all cohorts. Table shows median and (Quartile 1 – Quartile 3), ‡p<0.05 compared to both other cohorts, *p<0.05 compared to the Tromsø cohort.

| Parameter | Groningen | | Tromsø | | Texas | |
|---|---|---|---|---|---|---|
| Latency to first mount or intromission | 15.1 | (3.2-8.0) | 18.2 | (7.1-43.3)‡ | 3.9 | (2.5-12.6) |
| Number of mounts | 6.0 | (2.5-10.0) | 5.0 | (2.5-12.0) | 20.0 | (6.8-27.3)‡ |
| Number of intromissions | 7.0 | (5.0-10.5) | 13.0 | (9.0-16.5)‡ | 7.0 | (6.3-9.0) |
| Intromission ratio | 0.53 | (0.42-0.71) | 0.70 | (0.55-0.81)‡ | 0.28 | (0.19-0.54) |
| Number of mount bouts | 9.0 | (6.5-15.0)‡ | 18.0 | (12.0-23.5) | 20.5 | (13.8-28.0) |
| Mounts per mount bout | 0.66 | (0.33-0.85) | 0.31 | (0.20-0.53)‡ | 0.84 | (0.53-1.18) |
| Intromissions per mount bout | 0.77 | (0.53-0.83) | 0.76 | (0.61-0.87) | 0.37 | (0.27-0.63)‡ |
| Mount bout duration | 3.1 | (1.5-5.0)* | 1.2 | (0.7-1.7) | 1.7 | (1.0-3.5) |
| Time out duration | 34.8 | (24.1-45.6)* | 13.14 | (10.5-18.9) | 20.9 | (16.3-24.9) |
| Inter-intromission interval | 49.5 | (32.0-80.1) | 17.7 | (14.6-29.2)‡ | 56.2 | (28.7-99.4) |
| Latency to ejaculation | 411 | (208-614) | 266 | (166-439) | 528 | (262-728) |
| Post-ejaculatory interval | 270 | (240-351) | 267 | (236-304) | 352 | (321-386)‡ |

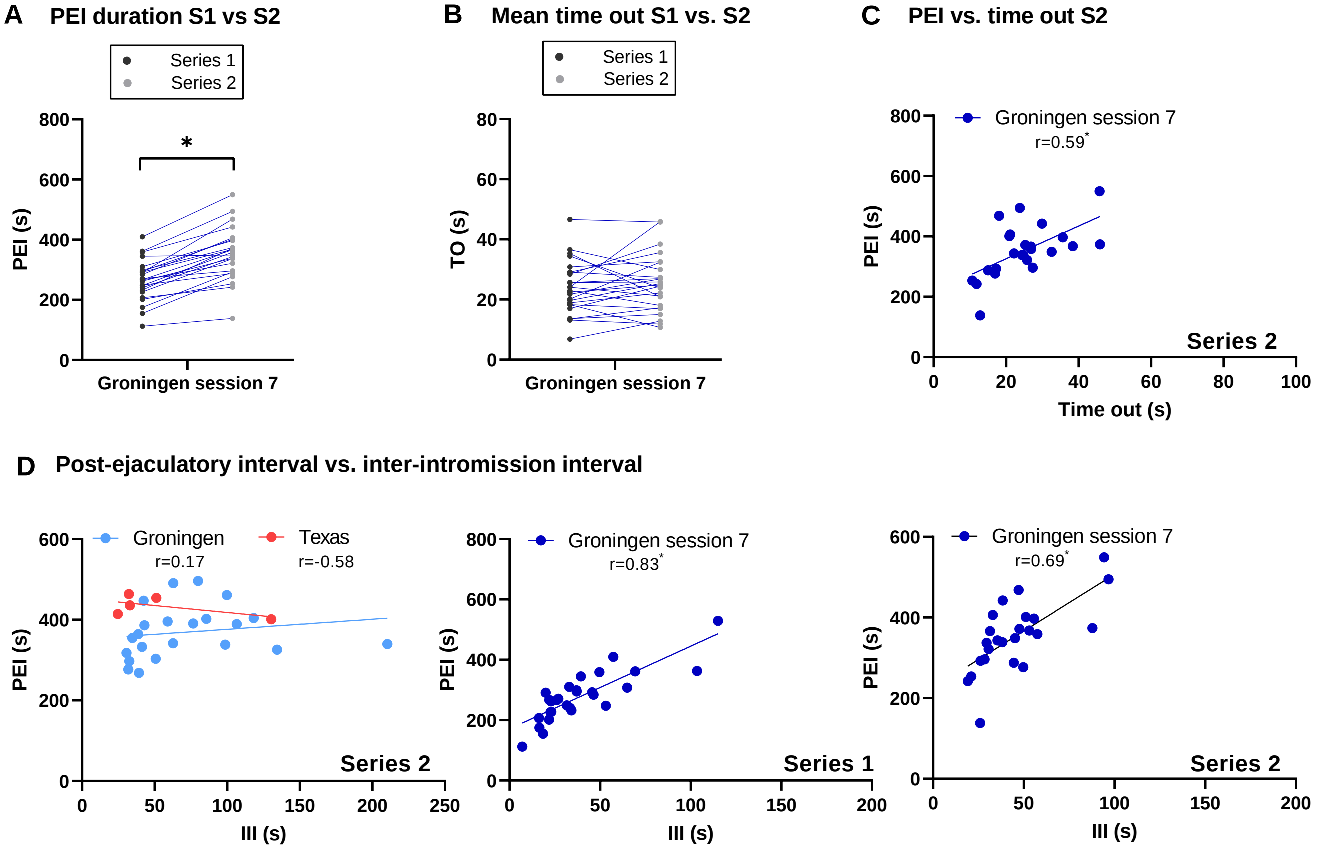


**Supplementary figure 1** Post-ejaculatory interval, time out, and inter-intromission interval over series in Groningen copulation session 7. **A** Post-ejaculatory interval duration in ejaculation series 1 compared to ejaculation series 2 within the same animals from the Groningen cohort in copulation session 7 (n=24). **B** Mean time out duration in ejaculation series 1 compared to ejaculation series 2 within the same animals from the Groningen cohort in copulation session 7 (n=24). **C** Correlation of post-ejaculatory interval and mean time out duration for ejaculation series 2 in copulation session 7 of the Groningen cohort (n=24). **D** Correlation of post-ejaculatory interval and inter-intromission interval for ejaculation series 2 in the Groningen (n=25) and Texas (n=6) cohorts, and for ejaculation series 1 (n=28) and 2 (n=24) in copulation session 7 of the Groningen cohort. **All panels:** $^{*}p<0.05$

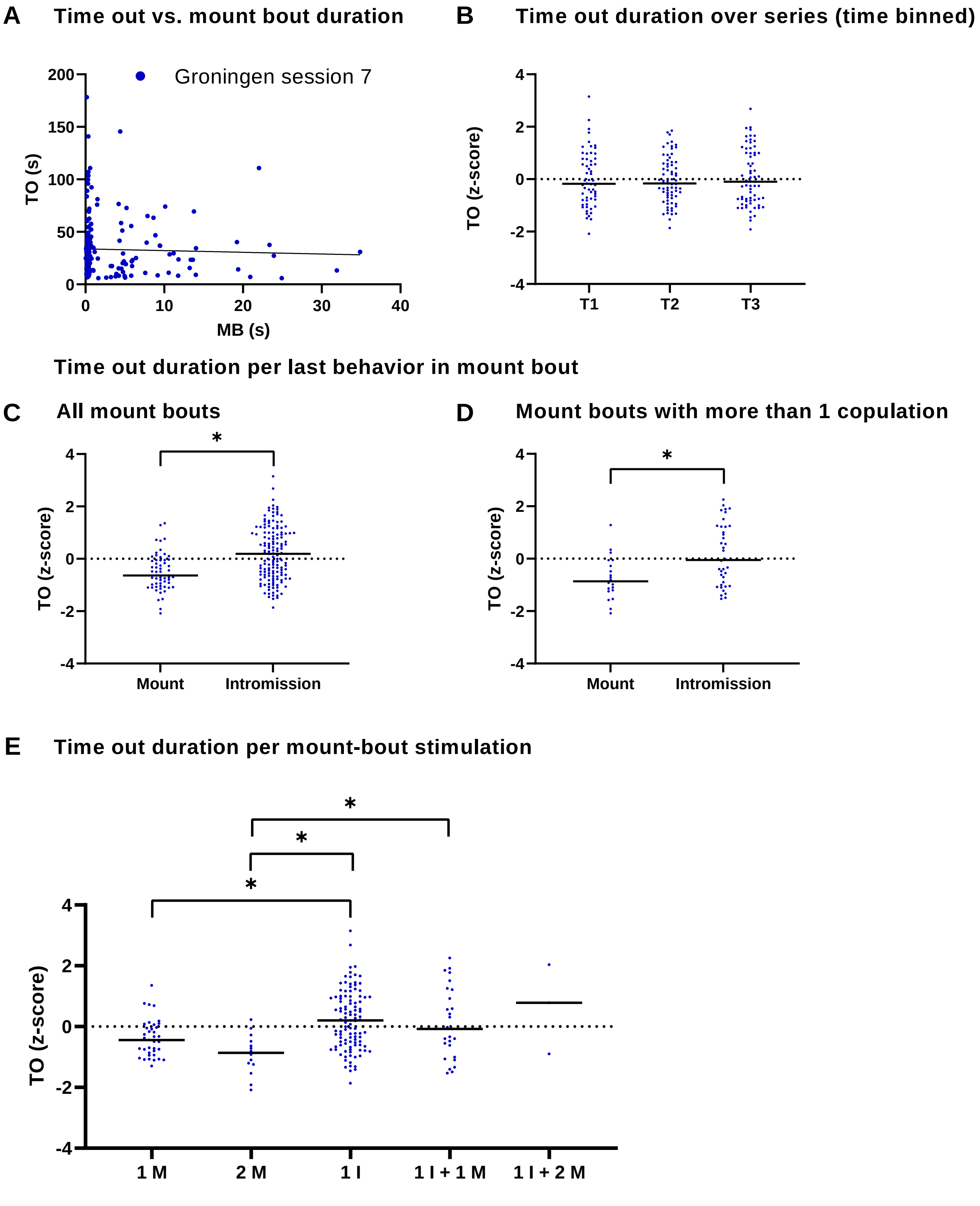


**Supplementary figure 2** Time out duration per mount bout characteristic for Groningen copulation session 7. **A** Correlation of individual mount bout duration with subsequent time out duration (n=202). **B** Z-scores of individual time out durations during the first, second, and third third of the ejaculation series (n=58; 71; 67). **C** Z-scores of individual time out duration after mount bouts with mount vs. intromission as last copulation (n=57; 145). **D** Z-scores of individual time out duration after mount bouts consisting of multiple copulations with mount vs. intromission as last copulation (n=20; 38). **E** Z-scores of individual time out durations after mount bouts with different total copulatory stimulation (n=37; 14; 107; 27; 3), M; mount, I; intromission. **All panels:** TO; time out, horizontal lines; median, $^{*}p<0.05$

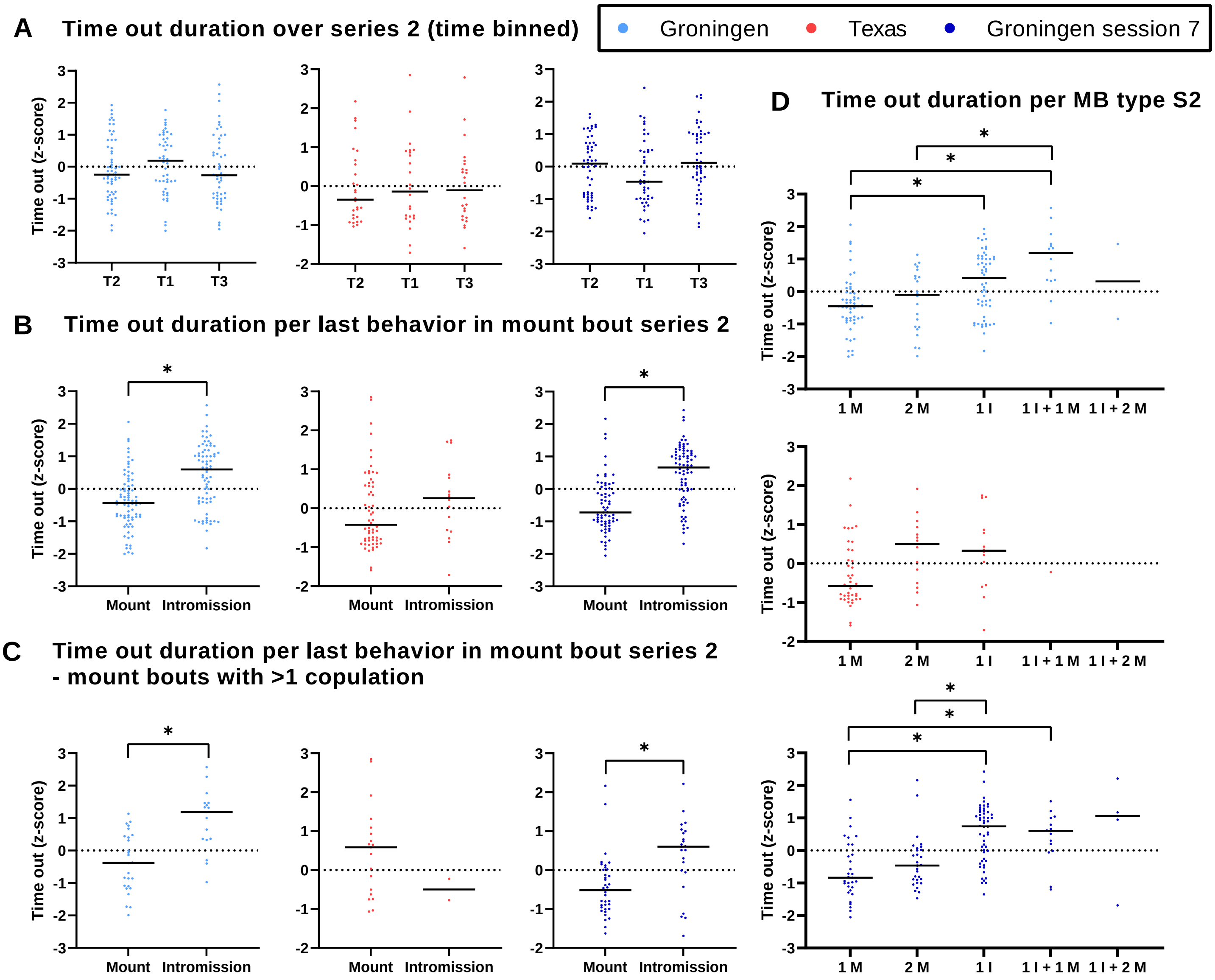


**Supplementary figure 3** Time out duration per mount bout characteristic for ejaculation series 2 in the Groningen and Texas cohorts, and Groningen copulation session 7. **A** Z-scores of individual time out durations during the first, second, and third third of the ejaculation series (GR n=46; 57; 50), TX n=24; 28; 24, GR7 n=43; 52; 49)). **B** Z-scores of individual time out duration after mount bouts with mount vs. intromission as last copulation (GR n=79; 74, TX n=60; 16, GR7 n=67; 77). **C** Z-scores of individual time out duration after mount bouts consisting of multiple copulations with mount vs. intromission as last copulation (GR n=26; 17, TX n=19; 2, GR7 n=36; 23). **D** Z-scores of individual time out durations after mount bouts with different total copulatory stimulation (GR n=53; 22; 57; 15; 2, TX n=41; 14; 14; 1, GR7 n=31; 29; 54; 15; 4), M; mount, I; intromission. **All panels:** horizontal lines; median, $^{*}p<0.05$

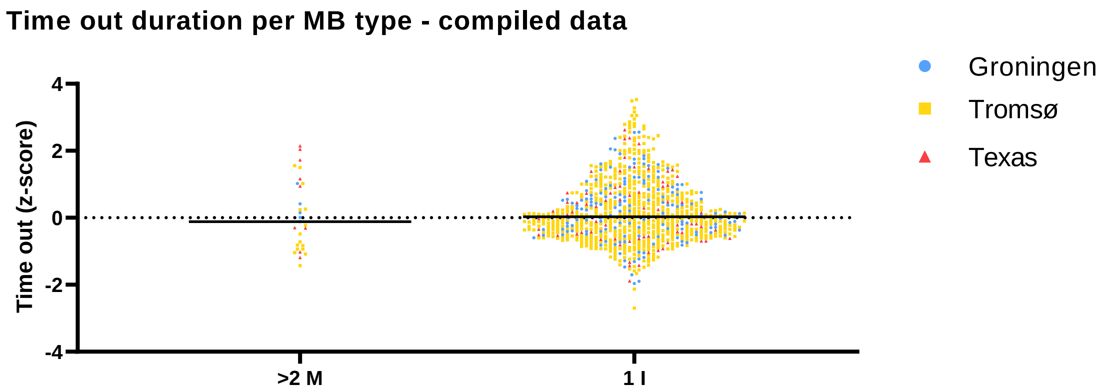


**Supplementary figure 4** Compiled data from Groningen, Tromsø, and Texas cohorts, shows no difference in time out duration following mount bouts of more than 2 mounts vs. mount bouts of 1 intromission, GR n=152; 5, TR n=640; 15, TX n=70, 9, horizontal lines; median.

**Post-ejaculatory interval vs. number of intromissions**

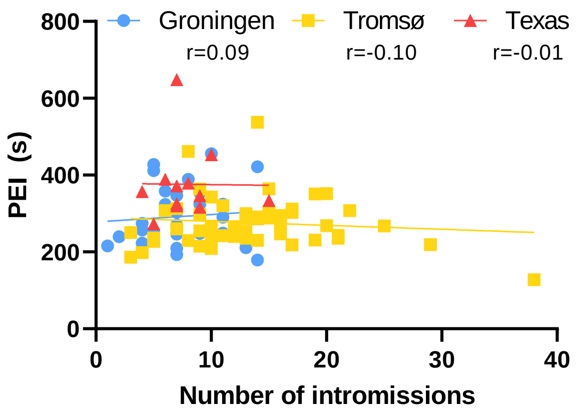


**Supplementary figure 5** Correlation of post-ejaculatory interval and number of intromissions for ejaculation series 1 for Groningen, Tromsø, and Texas cohorts. n=29; 53; 12
